# Bacteriophages inhibit biofilms formed by multi-drug resistant bacteria isolated from septic wounds

**DOI:** 10.1101/863076

**Authors:** Roja Rani Pallavali, Vijaya Lakshmi Degati, Vijaya Raghava Prasad Durbaka

## Abstract

Globally, indiscriminate use of antibiotics contributed to the development of antibiotic resistance by the majority of microbial pathogens. As an alternative to antibiotics, using bacteriophages as antibiofilm agents to tackle multi-drug resistant bacteria has gained importance in recent years. In the present study, we explored the ability of bacteriophages to inhibit biofilm formation under various conditions. Under dynamic condition (DR), wherein the medium is a renewal for every 12 h amount of biomass produced (0.74 ± 0.039), log_10_ CFU count (6.3 ± 0.55) was highest when compared to other physical conditions tested. Biomass of biofilms produced by *Staphylococcus aureus, Pseudomonas aeruginosa, Klebsiella pneumoniae*, and *Escherichia coli* drastically reduced when incubated for 2 or 4 h with bacteriophages vB_SAnS_SADP1, vB_PAnP_PADP4, vB_KPnM_KPDP1, and vB_ECnM_ECDP3 respectively at the time points tested (24, 48 and 72 h). Among the phages, vB_ECnM_ECDP3 effectively inhibited the biomass of biofilm when incubated for 2 h (0.35 ± 0.04, (44 %) (p < 0.0001) or 4 h (0.17 ± 0.015, (21.5%) (p<0.0001). Bacteriophages of *E. coli* (vB_ECnM_ECDP3) *P. aeruginosa* (vB_PAnP_PADP4), *K. pneumoniae* (vB_KPnM_KPDP1) and *S. aureus* (vB_SAnS_SADP1) also significantly inhibited the biomass of biofilm formation as evidenced by Scanning Electron Microscopy and Confocal laser scanning microscopy.

## Introduction

Most multi-drug resistant microorganisms develop biofilms on the sites of wounds, posing difficulties in treatment with available antibiotics and thereby causing morbidity or mortality in infected patients (Karthik, 2014; Jin et al., 2019; Yuan et al., 2019). Biofilms are formed by the association of a variety of complex extracellular components such as polysaccharides, proteins, lipids, toxins, metabolites and DNA (Chan and Abedon, 2015; Sharma et al., 2016). Biofilms always tend to adhere to reliable support for maintaining structural integrity (Deshpande Kaistha and Kaistha, 2017; Liu and Yang, 2019). Pathogenic bacterial biofilms are associated with both acute and chronic infectious sites and provide protection to the bacteria from antimicrobial agents; since the drugs cannot penetrate this structural barrier (Costerton et al., 2003; Deshpande Kaistha and Kaistha, 2017). Therefore, pathogenic bacteria that form biofilms develop antibiotic resistance and are often found to be very difficult to be eliminated by the conventional antibiotics (Nouraldin et al., 2016; Oliveira et al., 2017; Fang et al., 2018).

Contrary to the planktonic bacterial cells, pathogenic bacterial communities in the extracellular matrix usually exhibit distinct features such as:: a) intercellular signals between the community (quorum sensing) (Sharma et al., 2016), which regulates the maturation and detachment of the biofilms; b) activation of secondary messengers, which plays a role in the formation of biofilms, flagellar movements and production of extracellular polysaccharides (Hoggarth et al., 2019) and c) expression of DNA-binding proteins, amyloid and amyloid-like proteins, Biofilm Associated Proteins (Bap), and critical proteins which play a significant role in the establishment of the matrix association for the bacteria to encase in biofilms (Zhu et al., 2002; Le et al., 2013; Gondil and Chhibber, 2018; Schiffer et al., 2019). Formation of biofilms depends on both internal and external factors such as moist surface, energy source at the wound site, type of bacterial interaction, availability of receptors on bacteria for its attachment, temperature, pH and other factors (Merckoll et al., 2009; Bessa et al., 2015; Zhang et al., 2018). Hence, interventions at any of the unique features of pathogenic bacterial communities using unconventional bacterial clearing agents (bacteriophages) might provide newer strategies for the management of pathogenic diseases (Alharbi and Zayed, 2014; Vinodkumar et al.; Kumari et al., 2009; Dias et al., 2013).

Bacteriophage based therapy has been in practice for several years (Summers, 2001; Sillankorva et al., 2008; Burrowes et al., 2011; Golkar et al., 2014; Dalmasso et al., 2016). Bacteriophages also have been used as therapeutics for the treatment of bacterial diseases in plants, animals, as well as in humans (Delfan et al., 2012; Azizian et al., 2013; Zaczek et al., 2015). Bacteriophages when delivered at the infection site can replicate to produce a large number of active progeny, which in turn produce elution factors and degrading enzymes that target the polysaccharide components of the host bacteria, leading to bacterial lysis (Ghannad and Mohammadi, 2012; Vieira et al., 2012; Nouraldin et al., 2016). Though many studies have shown bacteriophage mediated mechanism of biofilm degradation, very few studies have used biofilms formed by clinical isolates (Pires et al., 2011; Danis-Wlodarczyk et al., 2015; Deshpande Kaistha and Kaistha, 2017; Oliveira et al., 2017; Ribeiro et al., 2018). Hence, there is an urgent need to identify bacteriophages that can act on the biofilms formed by bacteria isolated from wounds.

In this study, we investigated the ability of bacteriophages to degrade biofilms formed by single species pathogenic bacteria under different physical conditions (static and dynamic with and without renewal of media for every 12 h of incubation). Single species bacteria used in this study could form colony-forming units and biofilms with the maximum biomasses under the DR condition, i.e., dynamic condition with the renewal of media for every 12 h of incubation. Scanning and confocal electron microscopic studies indicated the inhibitory effect of respective bacteriophages on gram-positive (*S. aureus*) and gram-negative (*P. aeruginosa, E. coli*, and *K. pneumoniae*) bacterial biofilm biomasses. Our observations provide information that supports the ongoing efforts for establishing bacteriophages as antibiofilm agents against multi-drug resistant bacteria.

## Results

### Characterization of bacteriophages

For practical purposes, the phages were named based on Ackerman classification, which relies on the morphology of tail features (Ackermann, 2001; Ackermann et al., 2015). Morphological features of phages vB_PAnP_PADP4, vB_ECnM_ECDP3, vB_KPnM_KPDP1, and vB_SAnS_SADP1 as observed by Transmission Electron Microscopy are shown in **Figures 1A, 1B, 1C and 1D** respectively. The one-step growth curve of vB_PAnP_PADP4, vB_ECnM_ECDP3, vB_KPnM_KPDP1, and vB_SAnS_SADP1 are shown in Figures **1E**, **1F, 1G and 1H** respectively. The morphological features, latent period, and the burst size obtained basing on TEM, and the one-step growth curve is presented in Table 1.

**Figure 1.**
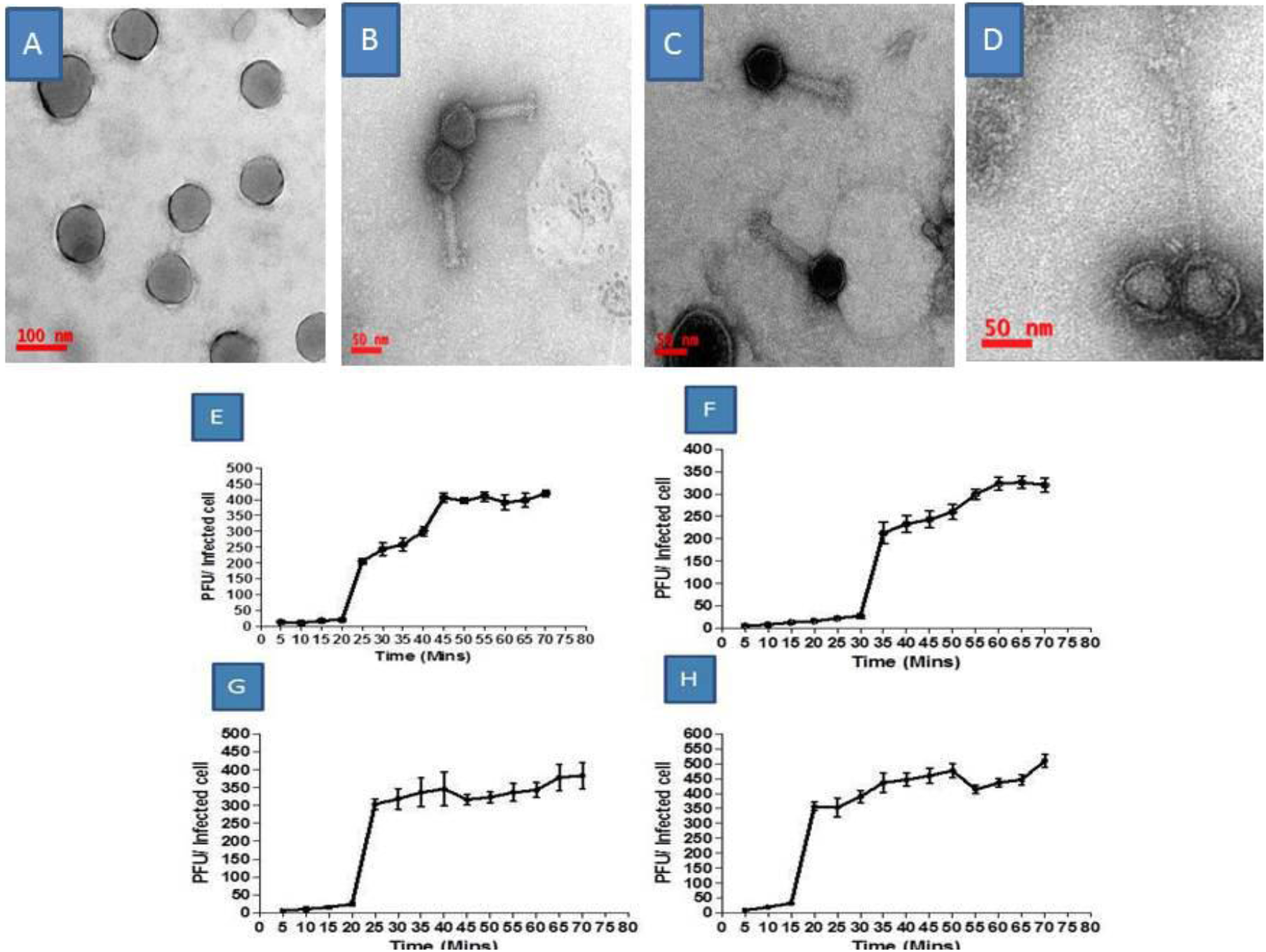
Morphology and one-step growth curve of bacteriophages. Phages are negatively stained with the 0.5% Uranyl acetate and visualized with scale bars represented A. vB_PAnP_PADP4 (100 nm), B. vB_ECnM_ECDP3 (50 nm), C. vB_KPnM_KPDP1 (50 nm), and D. vB_SAnS_SADP1 (50 nm) at 80,000 X magnification with transmission electron microscopy. Latent period and burst size of phages as follows, E. vB_PAnP_PADP4 (20 min, 102 / bacterial cell), F. vB _SAnS_SADP1 (30 min, 126/bacterial cell, G. vB _KPnM_KPDP1 (20 min, 76 / bacterial cell) and H. vB_ECnM_ECDP3 (15 min, 144 / bacterial cell).

**TABLE 1.** Features of bacteriophages used in this study

| Host Bacteria → | *P. aeruginosa* | *E. coli* | *K. pneumoniae* | *S. aureus* |
| --- | --- | --- | --- | --- |
| Phage | PA DP4 | EC DP3 | KP DP1 | SA DP1 |
| Name | vB_PAnP_PA DP4 | vB_ECnM_ECDP3 | vB_KPnM_KPDP1 | vB_SAnS_S ADP1 |
| Family | Podoviridae | Myoviridae | Myoviridae | Siphoviridae |
| Capsid (nm) | 55.5 | 65.5 | 68.2 | 65.9 |
| Tail length (nm) | 7.8 | 98.5 | 95.4 | 130 |
| Latent period | 20 | 15 | 20 | 30 |
| Burst Size | 102 | 144 | 76 | 126 |

### Stability of phages

The stability of vB_PAnP_PADP4, vB_ECnM_ECDP3, vB_KPnM_KPDP1, and vB_SAnS_SADP1 was investigated under different thermal and pH conditions. The percentage survival of phages were reduced by one order of magnitude after 60 and 90 min incubation periods at 40°C, 50°C, and 60°C (**Supplementary Figure 1A-D**), and the stability of phages at different range of pH showed that the phages were stable within the pH range 6 to 8 and showed 100 % activity (**Supplementary Figure 1E-H**). For all the phages tested, incubation at pH 5 and 9 caused 1 log decrease in the plaque-forming unit after 1 h, and incubation at pH 4 and 10 caused a 10-fold decrease in the plaque-forming units after 1h. Significant inactivation was observed at pH 2.0, and pH 12 for all the phages and some phages of vB_PAnP_PADP4, vB_ECnM_ECDP3, vB_KPnM_KPDP1, and vB_SAnS_SADP1 retain activity at pH 3 to pH 11 at different time intervals of 30, 60 and 90 min, the phages showed variations in their survival rate at various temperature and pH. The results suggest that extreme temperatures and pH conditions affect the stability of phages.

### Qualitative determination of biofilm formation of MDR-bacterial isolates

Previously isolated and screened MDR-bacteria from wound infections (Pallavali et al., 2017) were used for the qualitative determination of biofilm formation by various methods. **Supplementary Figure 2A, 2B, 2C, and 2D** show the representative biofilm formation on Congo red agar plates, 96 well culture plates, test tubes, and single-line well culture plates methods, respectively. The number of bacterial isolates that can form strong to moderate or weak biofilm is presented in **Figure 2A** basing on the intensity of color developed. The strength of color developed in a crystal violet staining test as a result of strong or moderate or weak biofilm formation is shown in **Figure 2B**. Among the total bacterial isolates used in this study, 28 strains of the four pathogens showed 64.26%, 24.99%, and 10.71% strong, moderate, and weak biofilm formation, respectively. Statistical comparisons among strong, moderate, and weak biofilm formers are presented in **supplementary Table 1.**

**Figure 2.**
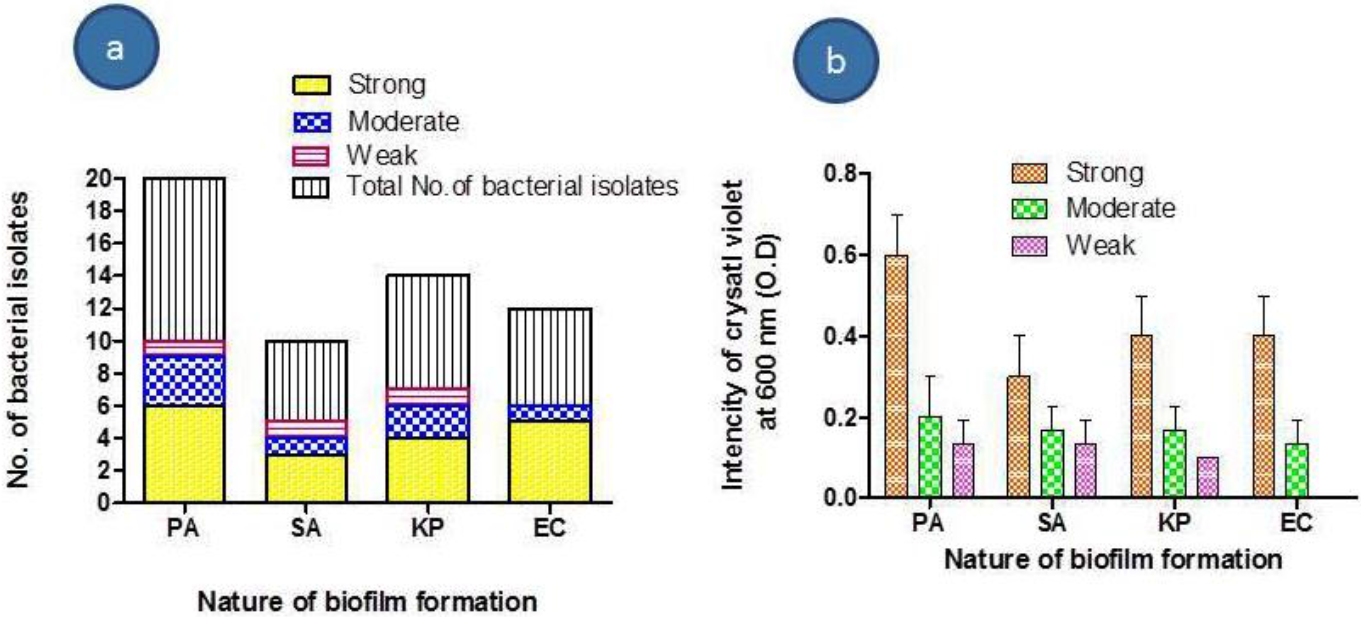
Type and Number of bacteria were used in the present study. a. The number of bacteria which were used for the present study and b. Based on O.D values differentiated into strong, moderate, and weak biofilm formers.

### CFU count and biomasses of the bacterial biofilms under various physical conditions

The dynamic condition with the renewal of media for every 12 h (DR) produced the highest CFU count and biomass than the remaining conditions, i.e., DNR and SR at different time intervals of 24, 48, 72, and 96 h **(Figure 3A).** *E. coli* and *K. pneumoniae* (6.3 ± 0.55) at 48 h of incubation under DR condition have the more log_10_ CFU count than the remaining conditions. Interestingly under SR condition at 48 h of incubation, *P. aeruginosa* (4.3± 0.66) has a higher CFU than the remaining bacteria, whereas under DNR condition at 24 h of incubation *S. aureus* (3.5 ± 0.30) had the highest CFU than the remaining bacteria. In almost all cases, after 72 h of incubation, reduction in CFU count was noticed, and in SR condition, *E. coli* and *K. pneumoniae* (1.6 ± 0.21) recorded the least CFU count than the remaining organisms used in the study. The mean log of CFU/mL of the single species bacteria at various conditions such as in SR, DNR, and DR showed the statistically significant at p < 0.05 by one way ANOVA (with p valve = 0.0089 and R square value = 0.4912) (**Supplementary Table 2)**.

**Figure 3.**
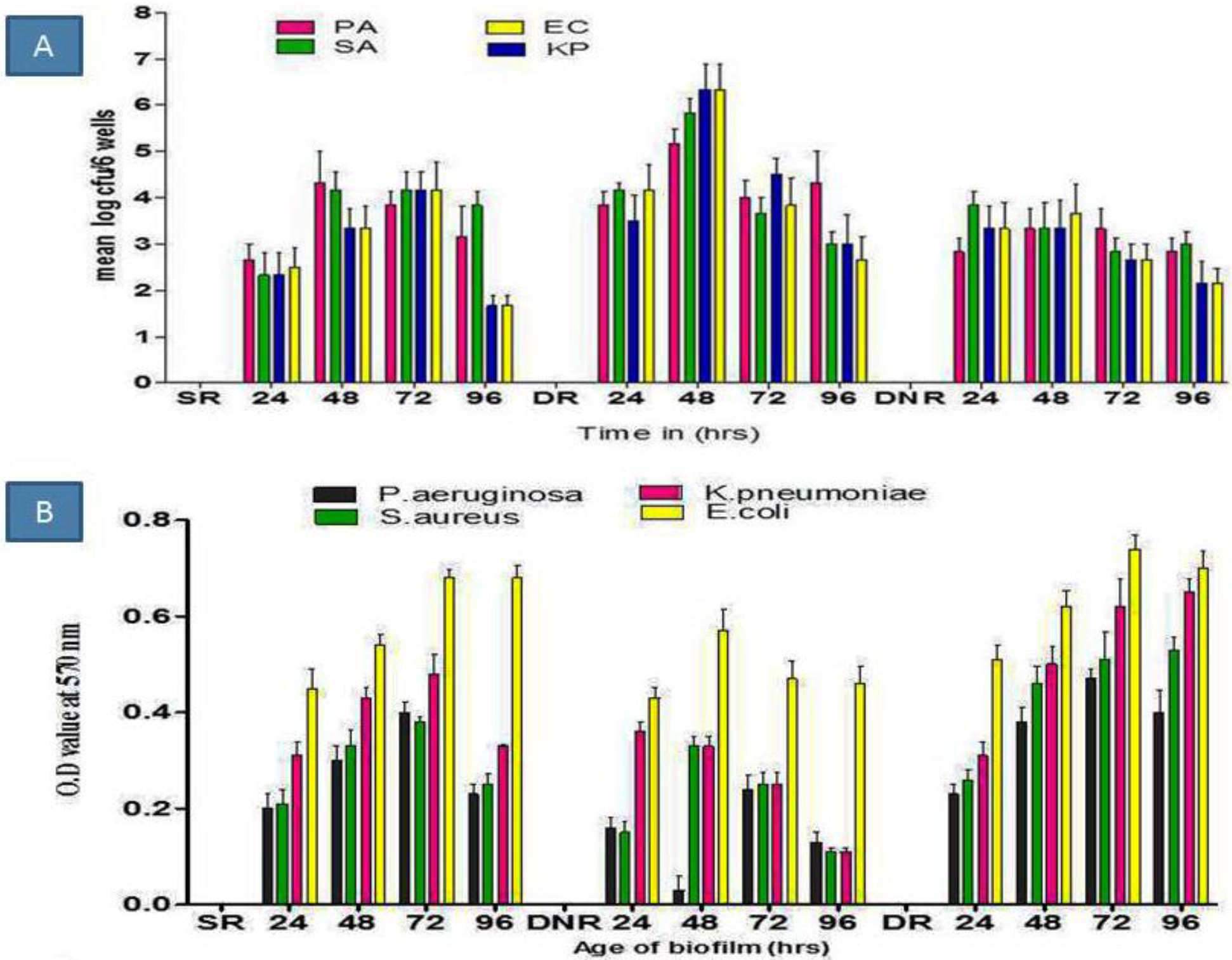
Determination of log CFU counts and biomasses of MDR-bacterial biofilms. A. Determination of log_10_ colony-forming units of biofilm of *P. aeruginosa, S. aureus, K. pneumoniae*, and *E. coli* at 24, 48, 72, and 96 h of incubation under static and dynamic with and without renewal of media for every 12 h of incubation (SR, DNR, and DR). The bars represent the mean values ± standard deviations (n = 6), each performed three times. Repeated ANOVA was done by using Graph pad Prism software. B. Determination of biofilm biomasses of *P. aeruginosa, S. aureus, K. pneumoniae*, and *E. coli* at 24, 48, 72, and 96 h of incubation under static and dynamic with and without renewal of media for every 12 h of incubation (SR, DNR, and DR). ANOVA Bonferroni’s multiple comparison tests were done by using Graph pad Prism software.

The biomasses of bacterial biofilms at various conditions (SR, DNR, and DR) and different incubation periods (24, 48, 76, and 92 h) are shown in **Figure 3B**. DR condition produced a high amount of biomass compared to other conditions. *E. coli* produced a high amount of biomass (0.74 ± 0.03) than the other bacteria in all the conditions. *P. aeruginosa* produced the least amount of biomass (0.13 ± 0.021) under DNR conditions than the remaining tested bacteria. Repeated measures ANOVA was performed to analyze the biomass of the single species biofilm biomasses with statistical significant at P < 0.05 (with p = 0.005, R square = 0.8257) (**Supplementary Table 3)**.

### Determination of biomasses of bacterial biofilm before and after respective phage treatment

The lytic activity on biofilms formed by *P. aeruginosa, E. coli, K. pneumoniae* and *S. aureus*, by vB_PAnP_PADP4, vB_ECnM_ECDP3, vB_KPnM_KPDP1, and vB_SAnS_SADP1 respectively were evaluated by quantification of biofilm biomasses using crystal violet assay. Biomasses of single-species biofilms with (for 2 or 4 h) or without phage treatment obtained after 24, 48, 76, and 92 h after treatment are presented in **Figure 4**. The amount of biomass production by *P. aeruginosa* was significantly reduced at all the time points tested and in all the physical conditions when treated with the respective phage for 2 (**Figure 4A**). The reduction was more pronounced when incubated with the phage for 4 hours. Biomass of *P. aeruginosa* at static conditions (SR) was high (1 ± 0.0) and the lytic activity of phage vB_PAnP_PADP4at 2 h (0.057 ± 0.03) (57%), and at 4 h (0.0238 ± 0.02) (23%), was observed which indicates a twofold reduction in biomass. In DNR conditions, the biomass obtained (0.86 ± 0.03) (86%) at 24 h was significantly reduced due to high lytic activity due to 2 h (0.39 ± 0.031) (31%) or 4 h (0.19 ± 0.026) (19%) treatment by phage vB_PAnP_PADP4. One way ANOVA was performed to analyze the biomass of 2, 4 h incubation with phage vB_PAnP_PADP4at 24, 48 72 and 96 h incubation of *P. aeruginosa* biomass is statistical significant at P < 0.05 (p <0.0001, R square value = 0.7891) and Bonferroni’s Multiple Comparison Test were performed to compare the 2, 4 h phage action at 24, 48, 76 and 92 h biofilm and is statistically significant at P < 0.005 **(Supplementary Table 4A)**.

**Figure 4.**
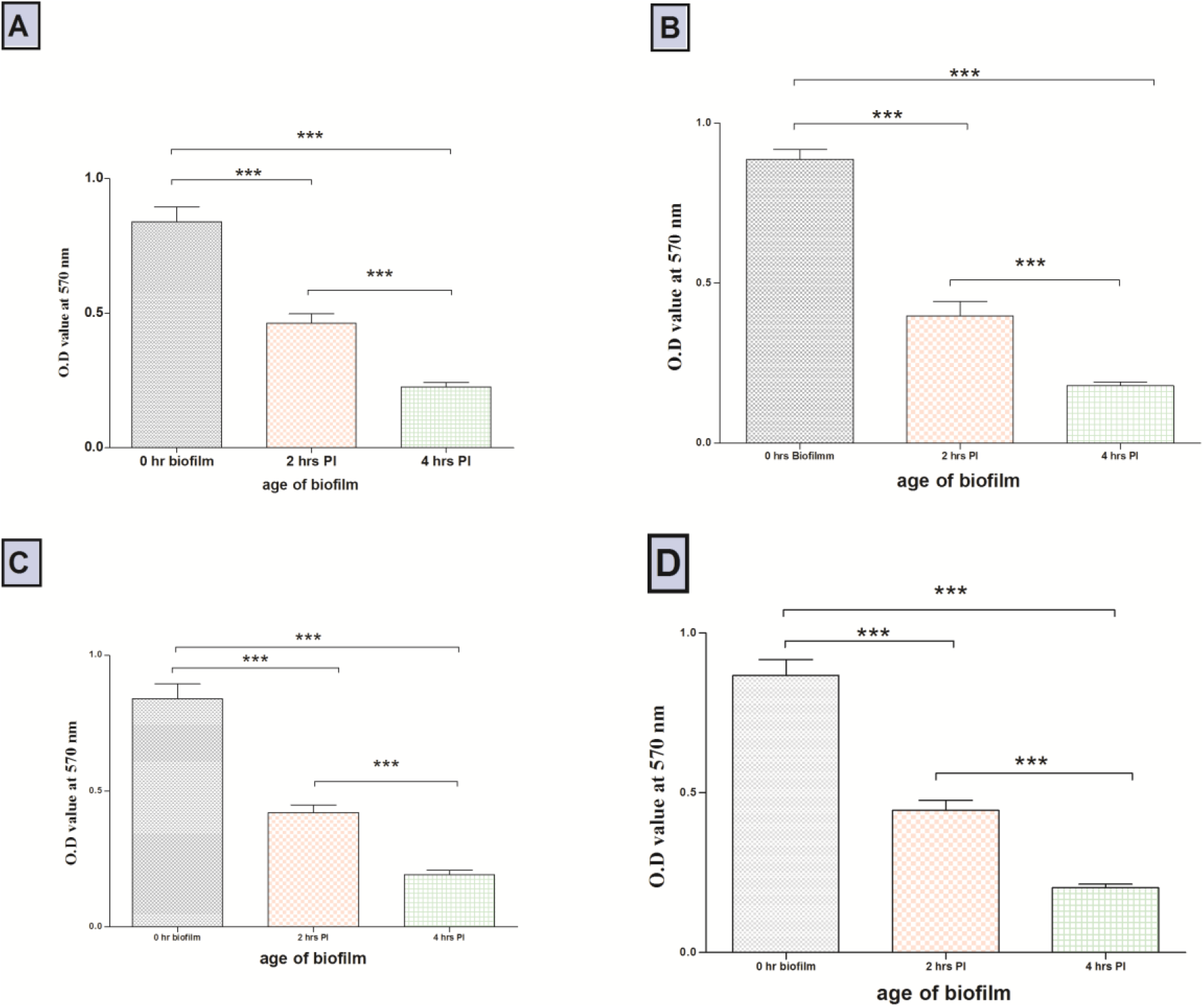
Bacteriophage lytic action towards their respective bacterial biofilm biomasses at 2 and 4 h incubation. Bacteriophages vB_PAnP_PADP4 (*P. aeruginosa)*, vB_SAnS_SADP1 *(S. aureus)*, vB_KPnM_KPDP1 (*K. pneumoniae)*, and vB _ECnM_ECDP3 (*E. coli*) inhibited the biomasses of their respective bacteria, at 2 and 4 h of phage incubation in SR, DNR and DR conditions. In all cases phage incubation (at 2 and 4 h) inhibited the biomasses nearly by 50% (2 h) to 80 % (4 h) at (P < 0.05). ANOVA and Bonferroni’s selective comparison tests were done by using Graph pad Prism software.

The biomass of the three other MDR-bacterial pathogens under SR, DNR, and DR conditions were also obtained (Figure 4B, C, and D). A high range of activity was noticed with *S. aureus* (Figure 4B) with the highest biomass at 48 and 72 h under SR condition, at 48 h (0.89 ± 0.03) under DNR and 48 h under DR conditions. Highest lytic activity was recorded at 4 h of incubation with SA DP1 (0.121 ± 0.08, 0.12 ± 0.051) (12%) and is statistically significant at P < 0.05 with p-value = <0.001, R square value = 0.8857 **(Supplementary Table 4B)**. For *K. pneumoniae* (Figure 4C), high biomass was observed under SR condition at 48 h and high lytic activity at 2 h (0.48 ± 0.03) (48%) and 4 h (0.21 ± 0.02) (21%) with phage KP DP1. Under DNR condition at 48 h (0.86 ± 0.038) whereas under DR maximum effect on biomass and lytic activity was observed at 48 h with similar activity noticed at 4 h (0.19 ± 0.02) (19%) with significant value of P < 0.05, R square value = 0.8293 **(Supplementary Table 4C)**. *E. coli* (Figure 4D)) bacteria showed the maximum extent of biomass at 48 h without phage application, whereas upon 2 h phage infection, the biomass was 0.47 ± 0.21 (47%) and it was 0.21 ± 0.02 (21%) when infected for 4 hours with EC DP3 **(Supplementary Table 4D)**.

### Biofilm eradication by bacteriophages was analyzed by SEM and CLSM

Since biofilms incubated for 2 or 4 h with phages showed decreased biomass and lysis, we used SEM and CLSM to gather more evidence on the morphological changes that occur during this process. *P. aeruginosa, S. aureus, K. pneumoniae*, and *E. coli* treated with respective phages under dynamic conditions **(Figures 6A, B, C, and D)** exhibited lesser biofilm formation when compared to untreated control. Treatment with respective phages resulted in a significant inhibition in biomass formation **(Figure 6 A1, B1, C1, and D1).** MDR-bacteria, when were grown under static conditions (**Figure 7**), showed biofilm formation to a more significant extent when compared to dynamic conditions **(Figure 7A, B, C, and D).** When these samples were subjected to lytic phages, destruction in the biofilm architecture with clear areas were observed in all the four cases **(Figure 7A1, B1, C1, and D1).**

**Figure 5.**
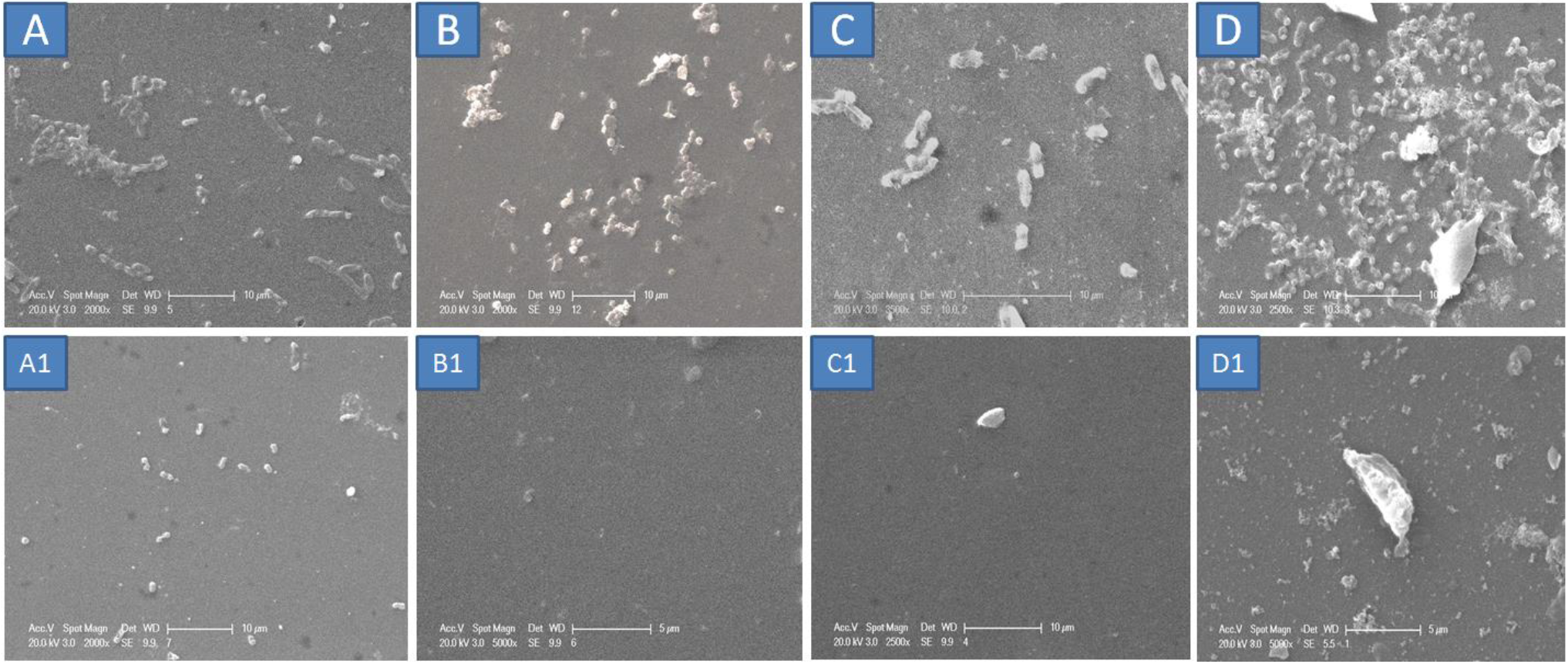
Scanning electron microscopic images of MDR-bacterial biofilms under dynamic condition treated with respective bacteriophages. Scanning electron micrographs of bacterial biofilms formed under dynamic conditions before and after the application of bacteriophages. A. *E. coli* A1 (vB_ECnM_ECDP3), B. *S. aureus*; B1 (vB_SAnS_SADP1), C. *K. pneumoniae*; C1 (vB_KPnM_KPDP1) and D. *P. aeruginosa* (vB_PAnP_PADP4) respectively and treated slides revealed less or little growth on the coverslip after 4 h of respective phage infections.

**Figure 6.**
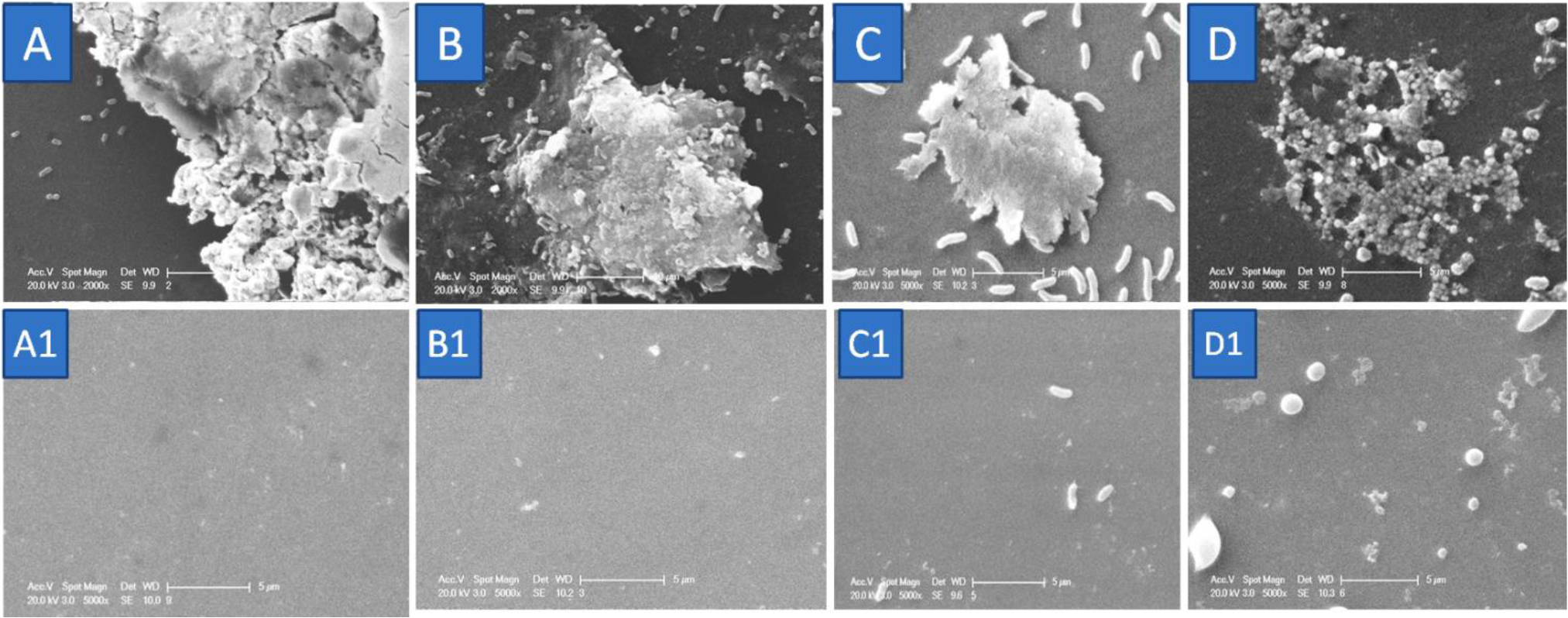
Scanning electron microscopic images of MDR-bacterial biofilm under static condition treated with respective bacteriophages. Scanning Electron microscopic images of *P.aeruginosa, E. coli, K. pneumoniae*, and *S. aureus* (A, B, C, and D) biofilm growing on the cover glass slip under static conditions. Control (only biofilm without respective bacteriophages) bacterial biofilms (A, B, C and D) forms the denced bacterial associations and phage treated groups A1 (vB_PAnP_PADP4), B1 (vB_ECnM_ECDP3), C1 (vB_KPnM_KPDP1) and D1 (vB_SAnS_SADP1) have no or little growth on the coverslip after 4 h of respective phage incubation.

**Figure 7.**
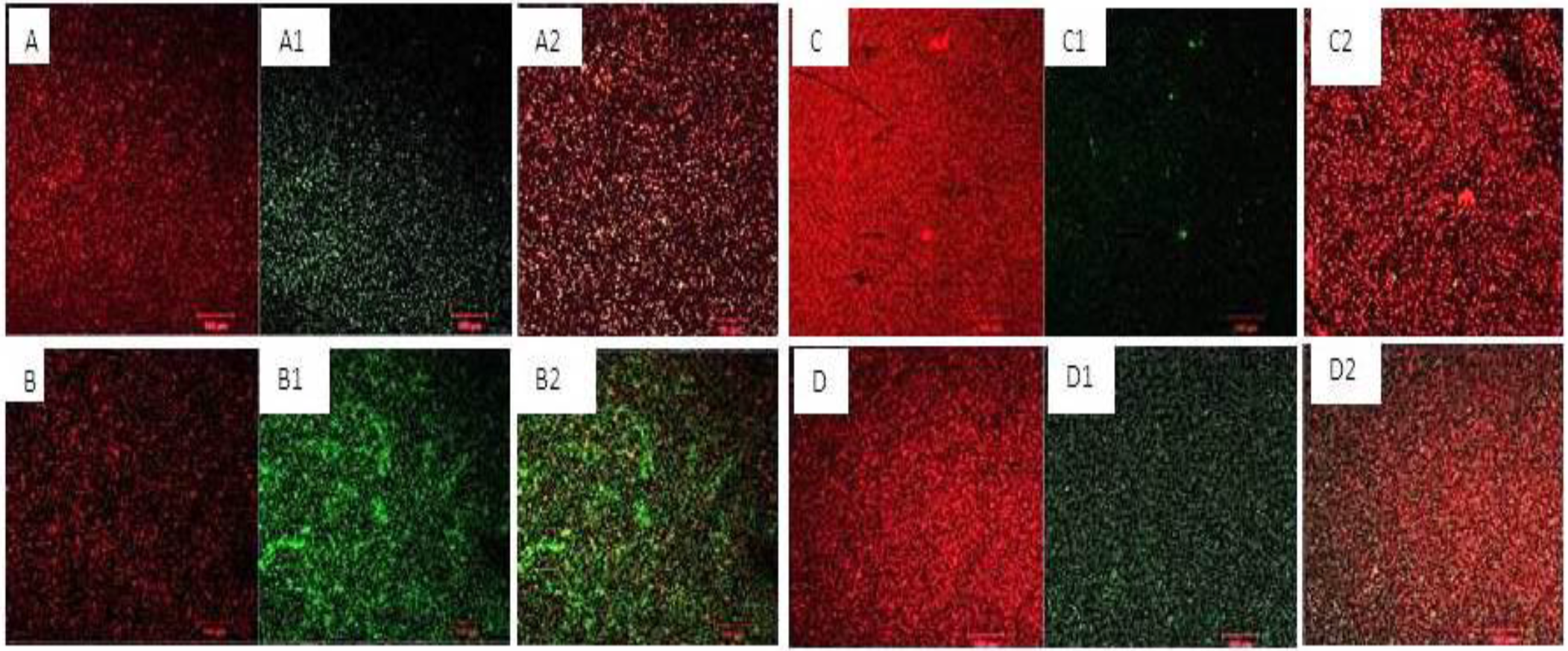
Determination of the lytic effect of bacteriophages on their specific bacterial biofilm by CLSM analysis. The biofilms of MDR-*P. aeruginosa, S. aureus, K. pneumoniae*, and *E. coli* were stained with SYTO^®^ 9 (green color indicates live cells) and propidium iodide (red color indicates dead cells). A, B, C, and D, the biofilms were treated with 4 h of phage incubation, A1, B1, C1, and D1 were treated with SM buffer (Control), and A2, B2, C2, and D2 were treated with 2 h of phage incubation. Red stained cells (dead cells) in the slide indicated that the lytic effect of bacteriophages on their respective bacteria. (Scale bars represented 20 μM-100 μM).

The ability of bacteriophages to inhibit biofilm formation and the associated structural changes were also evaluated by confocal microscopy. **(Figure 8A1, B1, C1 and D1)** The arbitrary numbers of dead cells (red-colored) in the biofilm were evident due to bacteriophage treatment for 2 h of incubation **(Figure 8A2, B2, C2 and D2)** and 4 h of incubation **(Figure 8A, B, C, and D).** These results prove that phages effectively inhibit the biomasses of biofilms *invitro*.

## MATERIALS AND METHODS

### Bacteria, Bacteriophages and growth conditions

MDR-bacterial isolates were isolated from patients suffering from burn wounds or post-operative wounds, or diabetic surgical wounds. Isolation of bacteria and bacteriophages, selection of phages against MDR-bacteria, and screening of multidrug-resistant bacteria were carried out as described previously (Pallavali et al., 2017). Briefly, pus swab samples from the human subjects were collected while dressing the wounds. Informed consent was obtained from the patient or the guardian in case the patient is a minor. *P. aeruginosa* yvu1 (Genbank: KY018605.1), *S. aureus* yvu2 (Genbank: KY496615.1), *K. pneumoniae* yvu3 (Genbank: KY496614.1) and *Escherichia coli* yvu4 were selected and grown on Luria agar (Himedia, Mumbai, India), at 37°C. Bacteriophages vB_PAnP_PADP4 for *P. aeruginosa*, vB_ECnM_ECDP3 for *E. coli*, vB_KPnM_KPDP1 for *K. pneumoniae* and phage vB_SAnS_SADP1 for *S. aureus* were isolated and characterized. The bacteriophages were stored in salts of magnesium buffer (5.8gL^-1^NaCl, 2gL^-1^ MgSO4.7H2O, 1M Tris HCl pH 7.5) at 4°C. Biofilms were grown in brain heart infusion broth media with 5% glucose at 37°C, under various conditions such as dynamic condition without renewal of media for every 12 h (DNR), dynamic condition with renewal of media for every 12 h (DR) and static condition with renewal of media for every 12 h (SR) for 24 h to 96 h. Growing bacteria detected biofilm formations on Congo red agar media, test tube adherence test, and microtiter plate methods. The biofilm staining before and after phage treatment on the coverslip was performed using Film Tracer™ LIVE/DEAD^®^ Biofilm viability kit (Molecular Probes, Life Technologies Ltd) according to the manufacturer’s instructions. This study was approved by the Institutional Ethics Committee (IEC) of Yogi Vemana University (IEC/YVU/DVRP dt 11/10/2014)

### Characterization of Bacteriophages

#### Transmission electron microscopy

Transmission electron microscopy (TEM) was carried out to determine the morphological features of bacteriophages used in this study. TEM analysis was carried out with FEI Tecnai G2 S-Twin (Hillsboro, Oregon, US). The isolated bacteriophage filtrate was passed through 0.45-micron filters (Hi-Media, Mumbai) and concentrated by centrifugation at 30,000 g for 60 min (Beckmann Coulter Benchtop centrifuge, USA) the pellet obtained was mixed with the 5 ml of SM buffer. 5 μL of phage filtrates were placed on formvar coated 200 x 200 copper grids. Excess phage filtrates were removed with filter paper from the edges of the grid. 5 μL of 0.5% uranyl acetate was then applied to the grids, and excess solution was immediately removed, and grids were air-dried. Samples were viewed with the FEI Tecnai G2 S-Twin Transmission Electron at an operating voltage at 80 KV (Sangha et al., 2014; Kwiatek et al., 2015; Stalin and Srinivasan, 2016).

#### Single-step growth curve of bacteriophages

One step or single-step growth curves were generated to determine the latent period and burst size of the bacteriophages as described (Cave et al., 1985; Skurnik et al., 2007). In brief, 50 mL of selected MDR-bacterial cultures were incubated to mid-exponential phase (A_600_ = 0.6), and cells were harvested by centrifugation at 10,000 g for 30 sec at 4°C. The pellets were resuspended in 0.5 mL of LB media and mixed with 0.5 mL of the phage filtrates having a plaque-forming unit (PFU) of 1×10^9^ PFU. This mixture was allowed to stand for 3 min at 37°C to facilitate phage adsorption on to the host cells. The mixture was then centrifuged at 13,000 g for 2 min to remove the free phage particles. The pellet was re-suspended in 100 mL of LB medium, and culture was incubated at 37°C with shaking at 150 rpm. Samples were taken after every 5 min up to 70 min and after centrifugation at 13,000 g for 1 min, subjected to determination of phage titer by double-layer agar method. The assay was performed in triplicates. The latent period was defined as the time interval between the adsorption and the beginning of the initial rise in the phage count. The burst size of respective phages was calculated as the ratio of the final phage titer to the initial count of infected bacterial cells during the latent period (Agudelo Suárez et al., 2002; Kwiatek et al., 2015).

#### Thermal and pH stability of phages

Thermal stability tests were performed according to the method described (Karumidze et al., 2013; Mishra et al., 2014; Piracha et al., 2014). Phage filtrates (1 × 10^9^ PFU) were taken in micro-centrifuge tubes and treated at different temperatures (37°C, 40°C, 50°C, 60°C and 70°C) for 0, 30, 60 and 90 min. Bacteriophage grown at 37°C was considered as control. After incubation, the double-layer agar method was performed for each treated sample to evaluate their lytic ability of phages (Al-Mola and Al-Yassari, 2010; Madsen et al., 2013; Mishra et al., 2014; Silva et al., 2014). For the pH stability assay, phage filtrates (1× 109 PFU/mL) was inoculated in a series of tubes containing SM buffer at pH 2.0, 3.0, 4.0, 5.0, 6.0, 7.0, 8.0, 9.0, 10.0 and 11.0 and incubated at 37°C for 4 h. Bacteriophage grown at pH 7.0 was considered as control. After incubation double-layer agar method was performed for each treated sample to evaluate their lytic ability of phages. All assays were performed in triplicates, and the results were tabulated.

### Qualitative determination of biofilm formation

#### Test tube method

The establishment of biofilm was achieved by the tube adherence test (Christensen et al., 1985). Briefly, 10 mL of brain heart infusion broth (Himedia, Mumbai) was inoculated with *P. aeruginosa or S. aureus or E. coli or K. pneumoniae* (10^6^ cells) and incubated for 24 h at 37°C with 130 rpm. After incubation, the medium was removed, and the tubes were washed twice with PBS (pH 7.4). The tubes were stained using 0.1% crystal violet and air-dried for 24 h. The presence of stained layer material adhered to the inner wall of the tube indicates biofilm formation.

#### Congo red agar method

Biofilm formation was also detected by Congo Red agar method (Christensen et al., 1985; Harper et al., 2014). *P. aeruginosa or S. aureus or E. coli* or *K. pneumoniae* from septic wounds were cultured on the BHI broth (Hi-Media, Mumbai) with the addition of 0.08% Congo red (Kmphasol, Mumbai) supplemented with 1% glucose. The bacterial isolates were streaked on BHI agar medium and incubated at 37°C for 24 to 48 h. After incubation, the plates were observed for phenotypic characterization, change in the color of the colony formation. Biofilm formers appear in black on Congo red agar, whereas non-biofilm producers appear red.

#### Microtiter plate method

Biofilm formation was investigated by a semi-quantitative method using 96 well flat bottom tissue culture plates (Sanchez et al., 2013; Cassat et al., 2014). Overnight bacterial suspensions made in BHI broth with 1% glucose and grown to mid-log phase (A600 = 0.1) (10^6^ CFU/mL). One hundred μL of bacterial suspension was then inoculated into each well and incubated overnight at 37°C for 48 h. After the incubation, the plates were gently aspirated and washed with 1X PBS (pH 7.4) followed by staining with 100 μL of 0.1% crystal violet (Qualigens, Mumbai) for 30 min at room temperature. Excess crystal violet was removed by gently washing with water.

In the present assay, the biofilms that were fixed using crystal violet were dissolved in 95% ethanol. The biofilm was quantified by measuring the absorbance of the supernatant at 570 nm. Biofilm assays were performed in triplicate for each bacterial strain tested, and their mean absorbance values were determined and distinguished them as strong (> 0.24), moderate (0.12 to 0.24), weak (0.05 to 0.12) and zero biofilm formers (< 0.05) based on their O.D values at 570 nm (Sanchez et al., 2013; Yadav et al., 2015; Zameer et al., 2016).

#### Biofilm formation in 24 well tissue culture plates

Overnight cultures of *P. aeruginosa or S. aureus or E. coli* or *K. pneumoniae* were diluted to 10^6^ CFU/mL into fresh BHI broth supplemented with 1% glucose (BHIG). 100 μL of each culture was diluted with 100 μL of BHIG and placed into each well of 24 well plates. Three different experimental conditions were applied:: a) Static condition (SR) with renewal of media for every 12 h without shaking; b) Dynamic condition (DNR) without renewal of media and shaking at 130 rpm; and c) Dynamic condition (DR) with renewal of media for every 12 h of incubation and shaking at 130 rpm. The development of biofilms was monitored after 24 to 96 h after incubation.

After incubation, the planktonic bacteria were washed off with PBS buffer (pH 7.4) subsequently four times, and biofilms were fixed with 200 μL of methanol for 15 min, followed by the addition of crystal violet (Qualigens, Mumbai) and incubated for 15 min. The wells were then washed with water and dried for 2 h at room temperature. 200 μL of ethanol (95%) was added to dissolve the stain. The absorbance of eluted stain was measured at 570 nm in a spectrophotometer (Bio-Rad Laboratories, Hercules, CA, USA). All the measurements were made with samples obtained from triplicate experiments (Sillankorva et al., 2010; Cassat et al., 2014; Abbatiello et al., 2018).

For the determination of colony-forming units, the adhered cells were collected by scratching the bottom of the well (n = 6) from each bacterial sample, with a sterile swab and subsequently suspended in 9 mL of PBS buffer by vigorous vortexing for 1 min. Serial dilutions of these suspensions were plated on brain heart infusion agar medium, and the colonies formed were counted and expressed as log_10_ CFU/mL (Abbatiello et al., 2018).

#### Determination of phage activity on biofilm biomass before and after phage treatment

100 μL of MDR-bacterial cultures (*P. aeruginosa or S. aureus or K. pneumonia* or *E. coli*) and 100 μL of the corresponding bacteriophages (10^9^ PFU) were added to each well. Control wells were loaded with 200 μL of SM buffer. The above sets of cultures were incubated under SR, DNR, and DR conditions for 2 or 4 h at 37°C for the analysis of cultures.

After incubation, the planktonic bacteria were removed by washing twice with PBS buffer. The total biomass attached to each well in 24 well tissue culture plates were measured by crystal violet assay. The wells were washed four times with PBS (pH 7.4), and then biofilms were fixed with 200 μL of methanol for 15 min. Methanol was removed, and each well was added 200 μL of crystal violet (1% v/v, Qualigens, Mumbai) and incubated for 15 min. The absorbance of eluted stain was measured at 570 nm in a spectrophotometer (Bio-Rad Laboratories, Hercules, CA, USA), and triplicates were maintained (Pires et al., 2011; Coulter et al., 2014; Fong et al., 2017).

#### Scanning electron microscopy

Biofilms were grown on borosilicate glass coverslips previously placed into the wells of a 24 well microtiter plate. Biofilms formed on the coverslips were incubated with 100 μL of respective bacteriophages for 4 h. After treatment, the coverslips were washed twice with PBS and dried in an incubator for 20 h at 37°C. The biofilms coated on glass slides were fixed with glutaraldehyde (2.5%) and dehydrated through a series of graded ethanol (30-100%) for 5 minutes. Further, the glass slides were sputtered with gold after critical point drying, and the aggregated biofilms were examined using Scanning electron microscopy (FEI, Tecnai G-2S Twin) (Ceri et al., 1999; Ribeiro et al., 2018).

#### Biofilm slide preparation for CLSM

24 h old biofilms of MDR-bacteria and their respective bacteriophage treated (2, 4 h) slides were stained with SYTO ^®^ stain and propidium iodide nucleic acid dyes. The following steps are involved in the staining process. Briefly, a working solution of fluorescent stains was prepared by adding 3 μL of SYTO^®^ nine stains and 3 μL of propidium iodide (PI) stain to 1 mL of filter-sterilized water. 200 μL of staining solution was deposited on a glass coverslip surface coated with biofilms of selected MDR-bacterial isolates treated with respective phages. After 15 min incubation at room temperature in the dark, samples were washed with sterile saline for removing the excess dye and rinsed with water from the base of the support material. In order to minimize the air contact and maintain constant sample moisture conditions, a coverslip was used on the specimen (González et al., 2017). Due to the bacteriolytic activity of phages leads to the disruption of bacterial biofilm integrity and allows the propidium dye and appears in red color and live bacteria present in the biofilm appears in green color. The massive amount of red color is due to the bacteriolytic activity of phages on their respective bacterial biofilms.

#### Confocal Laser Scanning Microscopy

MDR-bacterial biofilms and respective phage treated slides were subjected to CLSM to detect the effect of bacteriophages on the MDR-bacterial biofilms. The staining with FilmTracer™ LIVE/DEAD^®^ Biofilm viability kit (Molecular Probes, Life Technologies Ltd) was performed according to the instructions provided by the manufacturer (Ansari et al., 2015; Hu et al., 2015; González et al., 2017).

#### Statistical analysis

The data were analyzed using the graph pad prism software (Graph Pad Software lnc, La Jolla, CA, USA). All the values were expressed as mean SD, and a significant difference between variations denoted by p-valve <0.05 were estimated Dunn s and Bonferroni’s multiple comparison test using one-way ANOVA.

## Discussion

Biofilm formation is one of the essential survival strategies of many bacteria. This process is characterized by bacterial accumulation and wrapping in the extracellular matrix containing bacteriological societies (Sanchez et al., 2013; Bjarnsholt et al., 2018). In the case of either acute or chronic wound infections or septic wounds, pathogenic bacteria tend to form bacterial biofilms on the surface of the wounded sites, and these colonized bacteria show enhanced resistance to existing antibiotics (Merckoll et al., 2009; Alves et al., 2014). Antimicrobial agents become ineffective since the biofilms encase the bacteria and hinder the movement and infiltration (Lu and Collins, 2007; Burrowes et al., 2011; Sharma et al., 2016). The dual contributions of biofilms, i.e., allowing bacterial survival during starvation and preventing the action of antibiotics, pose a serious to health risk management in people with infectious diseases (Reisner et al., 2014; Gabisoniya et al., 2016).

In most scenarios wherein pathogenic bacteria become drug-resistant due to biofilm formation, alternative treatment strategies have gained importance (Nouraldin et al., 2016; Oliveira et al., 2017; Fang et al., 2018). Bacteriophages, which lyse bacteria, are increasingly being identified as alternatives to antibiotics (Dalmasso et al., 2016; Deshpande Kaistha and Kaistha, 2017). Though in the recent years, several studies have projected bacteriophages as potential antibiofilm agents, the mechanisms that underlie their lytic ability, nature of interactions between bacteriophage and biofilm-embedded bacteria are not well reported (Letkiewicz et al., 2010; Bolger-Munro et al., 2013; Abedon, 2015). In the present study, we explored the biofilm-forming ability, CFU count and biomass formation of four pathogenic MDR clinical bacterial isolates and their clearance by bacteriophages under various physical conditions (dynamic with and without renewal of media for every 12 h of incubation and static condition with renewal media for 12 h).

The bacteriophages isolated against the MDR–bacteria (*P. aeruginosa, S. aureus, K. pneumonia*, and *K. pneumoniae* were of sewage origin. Nomenclature of the bacteriophages as vB_SAnS_SADP1, vB_PAnP_PADP4, vB_KPnM_KPDP1, and vB_ECnM_ECDP3 which act on *P. aeruginosa, S. aureus, K. pneumonia*, and *K. pneumoniae* respectively was done as per standard protocols. The isolated bacteriophages which showed the highest host specificity, thermal and pH stability, less latent period, and high burst size were selected and were named based on their tail morphological studies by using Transmission Electron Microscopy. Bacteriophages against *P. aeruginosa* and *E.coli* were predominant than the remaining phages against the *S. aureus*, *K. pneumonia*. Isolation of bacteriophages against *K. pneumoniae* (Kumari et al., 2010), *E. coli* (Fan et al., 2012), *S. aureus* (Li and Zhang, 2014) *P. aeruginosa* (Piracha et al., 2014) was reported earlier. The methodology and the characterization of bacteriophages (latent period and burst size) followed in this study agrees with the earlier reports. The latent period / burst size for PA DP4, SADP1, KP DP1, and EC DP3 was found to be 20 min / 102, 30 min / 126, 20 min / 76 and 15 min / 144 respectively. These parameters for the four bacteriophages are in agreement with previous studies (Larcom and Thaker, 1977; Abedon et al., 2001; Al-Mola and Al-Yassari, 2010; Mateus et al., 2014; Eriksson et al., 2015).

Physiological factors such as temperature, pH play a crucial role in phage – bacterial interactions. Temperature is one of the essential external factors for the phage infectivity because it has a direct effect on the metabolic activities of bacterial cells. Results from this study show that the isolated phages were thermally active even at 40°C, 50°C, and 60°C and phages remaining viable up to 60°C after 90 min of incubation. The maximum infectivity was observed at 37°C, and the least infectivity was found at 60°C (Kęsik-Szeloch et al., 2013; Piracha et al., 2014; Wang et al., 2016).

We also observed that phages displayed maximum infectivity at pH 7, and their optimum range of pH was 6-8, which is similar to the observations made in earlier studies (Taj et al.; Elbreki et al., 2014; Silva et al., 2014). Thus, the isolation, characterization, and testing the lytic activity of bacteriophages isolated in this study at different temperatures and pH confirmed with standard protocols reported earlier.

According to the recent reports, it is increasingly projected that the phages can be used for the eradication of both oral planktonic and biofilms formed by *Actinomyces naeslundii, Aggregatibacter actinomycetemcomitans, Enterococcus faecalis, Fusobacterium nucleatum, Lactobacillus spp., Neisseria spp., Streptococcus spp.*, and *Veillonella spp* (Ceri et al., 1999; Sillankorva et al., 2010; Cornelissen et al., 2011; Khalifa et al., 2018; Shan et al., 2018). The use of a phage cocktail (Phage DRA88 and phage K) exhibited intense lytic activity against the biofilm formed by *S. aureus*. That the environmental bacteriophage, which extensively reduces the biofilm formation by *V. cholera*, is a causal organism of gastroenteritis from freshwater contamination (Naser et al., 2017). In this study, we report the lytic action of vB_PAnP_PADP4, vB_SAnS_SADP1, vB_ECnM_ECDP3, and vB_KPnM_KPDP1, which act on *P. aeruginosa, S. aureus, E. coli*, and *K. pneumoniae*, respectively. Our results add much more evidence to the possible application of bacteriophages for the treatment of wounds infected with pathogenic bacteria.

Bacteriophages inhibit the biomass of biofilms formed by corresponding host bacteria (Pires et al., 2011; Alves et al., 2014; Taha et al., 2018). Lytic bacteriophages eradicate the biofilms by releasing lytic enzymes (Sutherland et al., 2004; Lu and Collins, 2007; Azeredo and Sutherland, 2008). In this study, for evaluation of the lytic action of bacteriophages, two methodologies were employed viz measurement of biomass and enumeration of CFU. In the present study, single-species biofilms of *P. aeruginosa, S. aureus, K. pneumoniae, and E.coli* were exposed to bacteriophages VB_PAnP_PADP4, VB_SAnS_SADP1, VB_KPnM_KPDP1, and VB_ECnM_ECDP3, respectively. About 50 % to 80% inhibition of biomasses formation was observed in a time-dependent manner (2 to 4 hours). Such a reduction in biomass was reported (Webber and Hughes, 2017), wherein the maximum reduction was obtained within 2 h to 5 h, depending on the type of biofilm studied. These results suggest that usage of phages on single-species biofilms can effectively attach to the respective bacterial host cells and destroys the structural integrity of bacteriological populations in the biofilms, and can cause the disassociation of biofilms into individual non-adherent cells which are also subjected for lysis (Tait et al., 2002; Abedon, 2015).

In the current study, the effect of bacteriophages on single-species biofilms of MDR-bacteria that were isolated and characterized from septic wounds under various physical conditions were investigated. To the best of our knowledge, we, at this moment, for the first time, report the isolation of prevalent MDR-bacterial isolates from septic wounds comprising a mixture of gramnegative and gram-positive bacteria, which can form biofilms and can be successfully grown on coverslips. Further, we also show that isolated bacteriophages are effective in the elimination of biofilms. A previous study indicated that the eradication of biofilms requires both antibiotics and bacteriophages (Verma et al., 2010). Further, it is suggested that combinational therapy with bacteriophages along with DNase enzymes degrades the biofilm matrix efficiently (Hughes et al. 2010). However, our studies demonstrate that the biofilm can be eradicated solely by the bacteriophage itself, which is a significant development in the search for newer agents in place of antibiotics.

Previous studies that focused on biofilm eradication under different physical conditions such as static, dynamic renewal and dynamic non-renewal were reported (Sillankorva et al., 2010; Williams, 2013; Coulter et al., 2014; Sagar et al., 2017). Most of the studies concluded that dynamic conditions are more favorable than the other conditions to access the biofilms. Sillankorva et al. reported that the high phage concentration ØIBB-PF7A bacteriophage could be highly efficient in removing *P. fluorescence* biofilms within 4 h of incubation. The same group also demonstrated that the cell lysis starts faster under dynamic than that of the static conditions, interestingly it was noted that the total relative biofilm reduction was not significant during 4 h in dynamic biofilms as compared to static biofilms. In principle under static conditions, the released progeny will attach to the neighboring bacteria within the biofilms, whereas in case of dynamic conditions, the progeny of virus can target and lyse the entire span of bacterial colonies within the biofilms because of agitation in the experimental setup (Azeredo and Sutherland, 2008; Sillankorva et al., 2010; Azeredo and Sillankorva, 2018). We observed that *K. pneumoniae* and *E. coli* showed a significantly higher number of cells as compared to *P. aeruginosa and S. aureus* under DR condition at 48 h of incubation; Under SR condition, significant growth of MDR-bacterial isolates exhibited successful colonization These observations are in correlation with previous studies (Hughes et al.2010; Ribeiro et al., 2018). While in the case of single-species biofilm formed by *S. aureus* and treated with phage (vB_SAnS_SADP1), we noticed degraded biofilm with some single coccus spread on the coverslip.

The lytic activity of bacteriophages was examined and confirmed by the CLSM. Disruption of the cell membrane due to lysis leads to penetration of propidium iodide stain, whereas live bacterial cells that are not lysed stain with SYTO^®^9 appear in green color (Ansari et al., 2015; Hu et al., 2015; González et al., 2017). The application of phages on MDR-bacterial pathogens present in the single species biofilms showed decreased biomass to almost 80%.

In conclusion, we report the successful isolation of bacteriophages from sewage samples. Further, it is observed that bacterial biofilm formation by *P. aeruginosa, S. aureus, K. pneumoniae, and E.coli* is best achieved under the dynamic growth condition with media renewal. The same conditions were found to be the best for the maximum lytic activity of bacteriophages. The single species biofilms were effectively inhibited alone by specific lytic bacteriophages with a concomitant reduction in biomass and CFU. SEM and CLSM confirm the physical changes that occur during the bacteriophage mediated lysis. These findings support the possible use of bacteriophages for the development of alternatives to antibiotics for the treatment of wounds infected with MDR bacteria.

## Acknowledgments

The authors are very grateful to the Yogi Vemana University for its facilities. Authors gratefully acknowledge to Dr. Suresh Yenugu, Associate Professor, Department of Animal Biology, Hyderabad Central University, for providing laboratory space and necessary instructions to complete the biofilm analysis part of this work and Department of Nanotechnology and central lab facility for TEM, CLSM, and SEM, University of Hyderabad, Telangana. The author (PRR) gratefully acknowledges UGC for financial support in the form of JRF and SRF. This research did not receive any specific grant from funding agencies in the public, commercial, or not-for-profit sectors.

## Author contributions statement

RRP and VRPD conceived and designed the experiments. RRP performed the experiments, data collection, and analysis. RRP prepared figures, analysis of results, and tables. RRP drafted the manuscript. RRP, VLD, VRN, KKV analysis, interpretation of findings. RRP, VLD, VRN, KKV, VRPD read and revised the manuscript. All authors were involved in reviewing the manuscript and approval for publication.

---

**Supplementary Table. 1.** Statistical comparion of biofilm formation of used bacterial species in the present studies

| Table Analyzed | Type of Biofilm formation |
| --- | --- |
| **Repeated Measures ANOVA** |  |
| P value | < 0.0001 |
| P value summary | *** |
| Are means signif. different? (P < 0.05) | Yes |
| Number of groups | 4 |
| F | 33.71 |
| R square | 0.9183 |
| **Was the pairing significantly effective?** |  |
| R square | 0.1186 |
| F | 4.941 |
| P value | 0.0269 |
| P value summary | * |
| Is there significant matching? (P < 0.05) | Yes |

| ANOVA Table | SS | df | MS |  |  |
| --- | --- | --- | --- | --- | --- |
| Treatment (between columns) | 95.50 | 3 | 31.83 |  |  |
| Individual (between rows) | 14.00 | 3 | 4.667 |  |  |
| Residual (random) | 8.500 | 9 | 0.9444 |  |  |
| Total | 118.0 | 15 |  |  |  |
| Tukey's Multiple Comparison Test | Mean Diff. | q | Significant? P < 0.05? | Summary | 95% CI of diff |
| Strong vs Moderate | 2.750 | 5.659 | Yes | * | 0.6047 to 4.895 |
| Strong vs Weak | 3.750 | 7.717 | Yes | ** | 1.605 to 5.895 |
| Strong vs total bacterial isolates | -2.500 | 5.145 | Yes | * | -4.645 to -0.3547 |
| Moderate vs Weak | 1.000 | 2.058 | No | ns | -1.145 to 3.145 |
| Moderate vs total bacterial isolates | -5.250 | 10.80 | Yes | *** | -7.395 to -3.105 |
| Weak vs total bacterial isolates | -6.250 | 12.86 | Yes | *** | -8.395 to -4.105 |

**Supplementary Table. 2.**
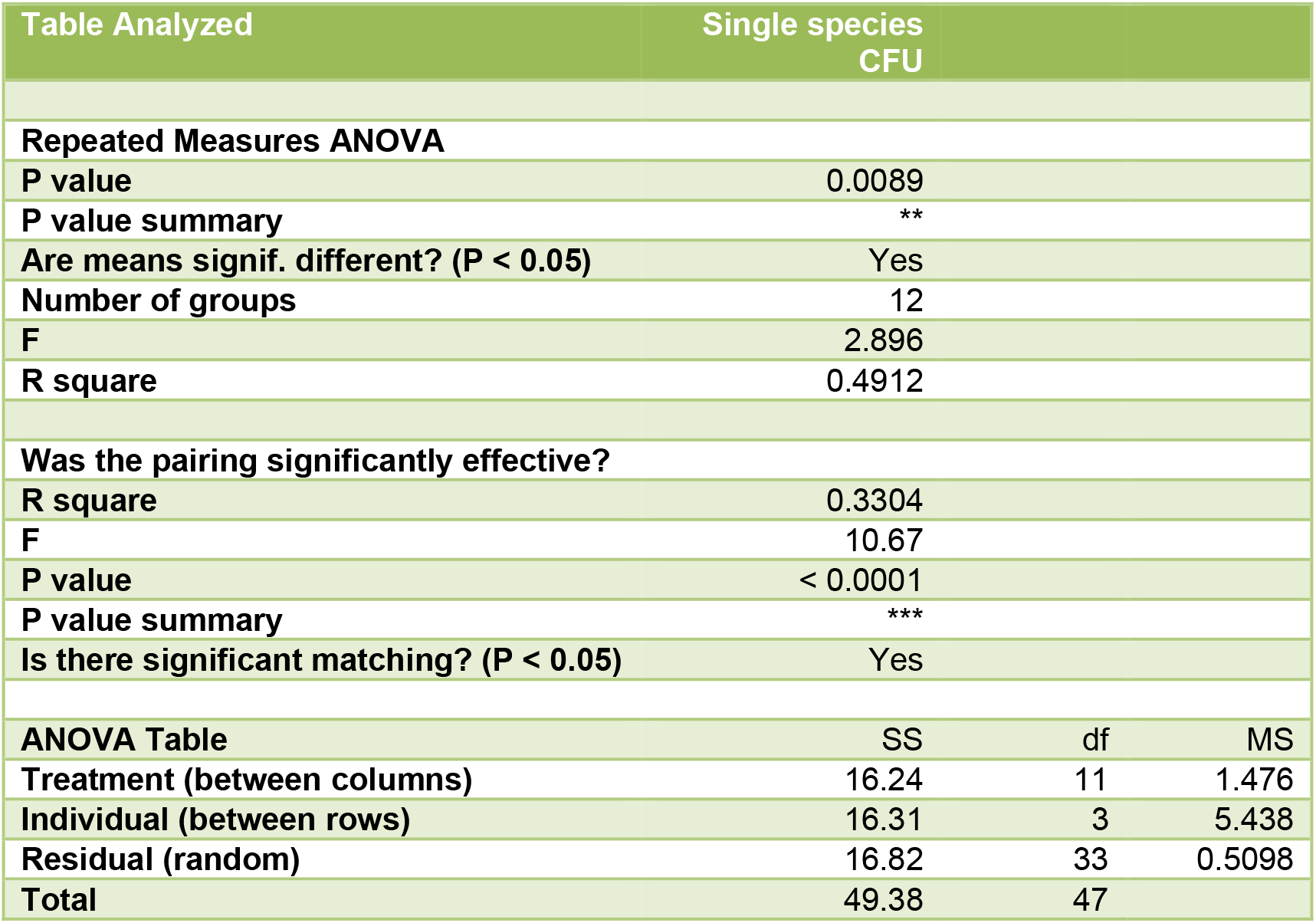
Statistical comparison of Mean log CFU count of bacteria under various conditions such SR, DNR, and DR

**Supplementary Table. 3.** Statistical comparison of Mean log Biomass amount of bacteria under various conditions such SR, DNR, and DR

| Table Analyzed | single species<br>biofilm biomass |  |  |
| --- | --- | --- | --- |
| Repeated Measures ANOVA |  |  |  |
| P value | < 0.0001 |  |  |
| P value summary | *** |  |  |
| Are means signif. different? (P < 0.05) | Yes |  |  |
| Number of groups | 12 |  |  |
| F | 14.21 |  |  |
| R square | 0.8257 |  |  |
| Was the pairing significantly effective? |  |  |  |
| R square | 0.1095 |  |  |
| F | 7.761 |  |  |
| P value | 0.0005 |  |  |
| P value summary | *** |  |  |
| Is there significant matching? (P < 0.05) | Yes |  |  |
| ANOVA Table | SS | df | MS |
| Treatment (between columns) | 1.052 | 11 | 0.09564 |
| Individual (between rows) | 0.1567 | 3 | 0.05222 |
| Residual (random) | 0.2221 | 33 | 0.006729 |
| Total | 1.431 | 47 |  |

**Supplementary Table. 4A.**
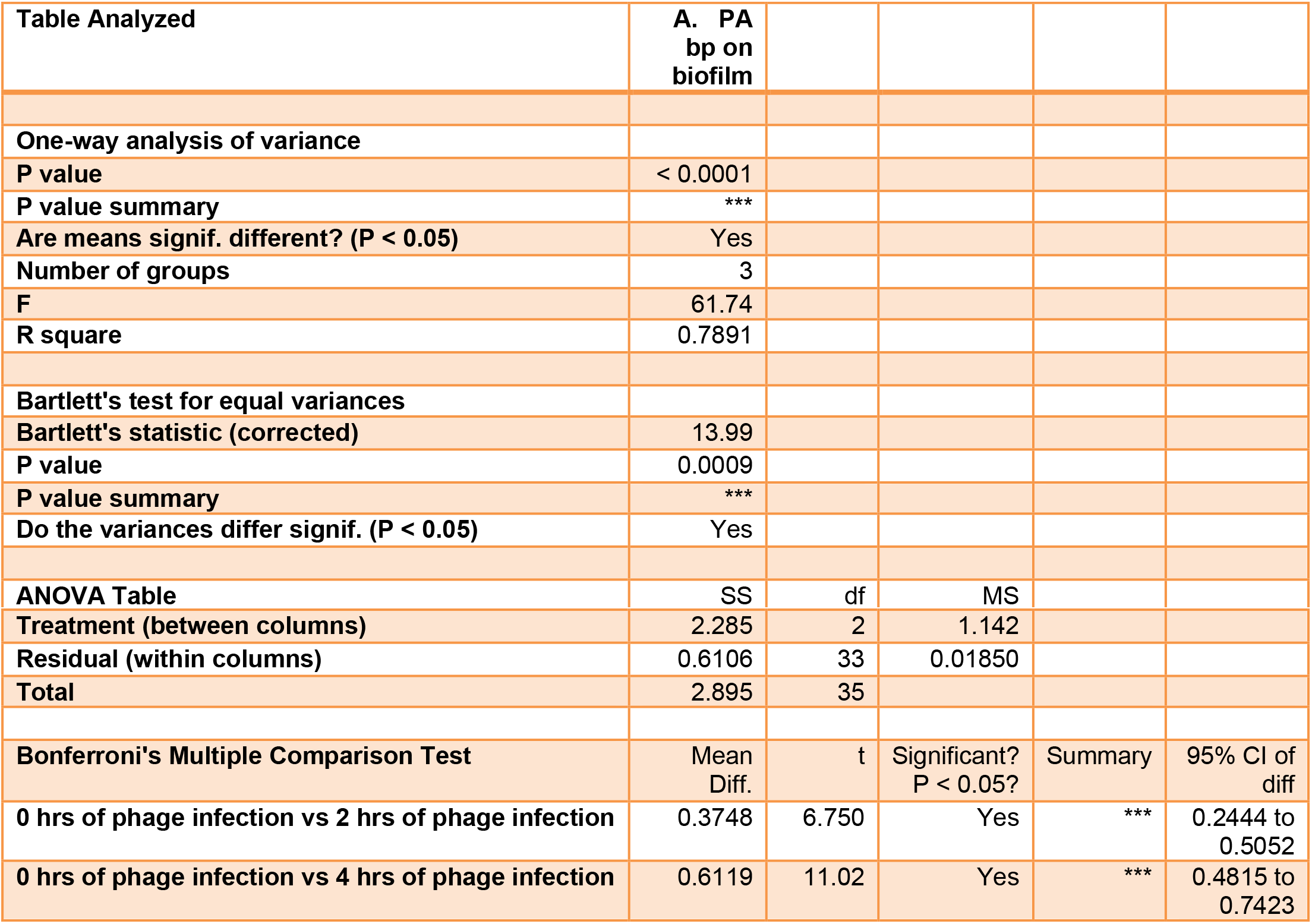
Statistical comparison of Mean log Biomass amount of *P. aeruginosa* with 2, 4 h of Phage vB_PAnP_PADP4 treatment under various conditions such SR, DNR, and DR.

| Table Analyzed | A. PA<br>bp on<br>biofilm |  |  |
| --- | --- | --- | --- |
| One-way analysis of variance |  |  |  |
| P value | < 0.0001 |  |  |
| P value summary | *** |  |  |
| Are means signif. different? (P < 0.05) | Yes |  |  |
| Number of groups | 3 |  |  |
| F | 61.74 |  |  |
| R square | 0.7891 |  |  |
| Bartlett's test for equal variances |  |  |  |
| Bartlett's statistic (corrected) | 13.99 |  |  |
| P value | 0.0009 |  |  |
| P value summary | *** |  |  |
| Do the variances differ signif. (P < 0.05) | Yes |  |  |
| ANOVA Table | SS | df | MS |

|  |  |  |  |  |  |
| --- | --- | --- | --- | --- | --- |
| **Treatment (between columns)** | 2.285 | 2 | 1.142 |  |  |
| **Residual (within columns)** | 0.6106 | 33 | 0.01850 |  |  |
| **Total** | 2.895 | 35 |  |  |  |
| **Bonferroni's Multiple Comparison Test** | Mean Diff. | t | Significant? P < 0.05? | Summary | 95% CI of diff |
| **0 hrs of phage infection vs 2 hrs of phage infection** | 0.3748 | 6.750 | Yes | *** | 0.2444 to 0.5052 |
| **0 hrs of phage infection vs 4 hrs of phage infection** | 0.6119 | 11.02 | Yes | *** | 0.4815 to 0.7423 |

**Supplementary Table. 4B.** Statistical comparison of Mean log Biomass amount of *Staphylococcus aureus* with 2, 4 h of Phage vB_SAnS_SADP1treatment under various conditions such SR, DNR, and DR.

| Table Analyzed | B. SA bp on biofilm |  |  |  |  |
| --- | --- | --- | --- | --- | --- |
| One-way analysis of variance |  |  |  |  |  |
| P value | < 0.0001 |  |  |  |  |
| P value summary | *** |  |  |  |  |
| Are means signif. different? (P < 0.05) | Yes |  |  |  |  |
| Number of groups | 3 |  |  |  |  |
| F | 127.9 |  |  |  |  |
| R square | 0.8857 |  |  |  |  |
| Bartlett's test for equal variances |  |  |  |  |  |
| Bartlett's statistic (corrected) | 14.57 |  |  |  |  |
| P value | 0.0007 |  |  |  |  |
| P value summary | *** |  |  |  |  |
| Do the variances differ signif. (P < 0.05) | Yes |  |  |  |  |
| ANOVA Table | SS | df | MS |  |  |
| Treatment (between columns) | 3.161 | 2 | 1.580 |  |  |
| Residual (within columns) | 0.4077 | 33 | 0.01236 |  |  |
| Total | 3.568 | 35 |  |  |  |
| Bonferroni's Multiple Comparison Test | Mean Diff. | t | Significant? P < 0.05? | Summary | 95% CI of diff |
| 0 hrs of phage infection vs 2 hrs of phage infection | 0.4903 | 10.81 | Yes | *** | 0.3838 to 0.5969 |
| 0 hrs of phage infection vs 4 hrs of phage infection | 0.7086 | 15.61 | Yes | *** | 0.6020 to 0.8151 |

**Supplementary Table. 4C.** Statistical comparison of Mean log Biomass amount of *Klebsiella pneumoniae* with 2, 4 h of Phage vB_KPnM_KPDP1treatment under various conditions such SR, DNR, and DR.

| Table Analyzed | C. KP bp on biofilm |  |  |  |  |
| --- | --- | --- | --- | --- | --- |
| One-way analysis of variance |  |  |  |  |  |
| P value | < 0.0001 |  |  |  |  |
| P value summary | *** |  |  |  |  |
| Are means signif. different? (P < 0.05) | Yes |  |  |  |  |
| Number of groups | 3 |  |  |  |  |
| F | 80.15 |  |  |  |  |
| R square | 0.8293 |  |  |  |  |
| Bartlett's test for equal variances |  |  |  |  |  |
| Bartlett's statistic (corrected) | 13.74 |  |  |  |  |
| P value | 0.0010 |  |  |  |  |
| P value summary | ** |  |  |  |  |
| Do the variances differ signif. (P < 0.05) | Yes |  |  |  |  |
| ANOVA Table | SS | df | MS |  |  |
| Treatment (between columns) | 2.586 | 2 | 1.293 |  |  |
| Residual (within columns) | 0.5324 | 33 | 0.01613 |  |  |
| Total | 3.119 | 35 |  |  |  |
| Bonferroni's Multiple Comparison Test | Mean Diff. | t | Significant? P < 0.05? | Summary | 95% CI of diff |
| 0 hrs of phage infection vs 2 hrs of phage infection | 0.4194 | 8.088 | Yes | *** | 0.2886 to 0.5502 |
| 0 hrs of phage infection vs 4 hrs of phage infection | 0.6472 | 12.48 | Yes | *** | 0.5164 to 0.7780 |
| 2 hrs of phage infection vs 4 hrs of phage infection | 0.2278 | 4.392 | Yes | *** | 0.09696 to 0.3585 |

**Supplementary Table. 4D.** Statistical comparison of Mean log Biomass amount of *Escherichia coli* with 2, 4 h of Phage vB_ECnM_ECDP3 treatment under various conditions such SR, DNR, and DR.

| Table Analyzed | D. EC bp<br>of Biofilm |  |  |  |  |
| --- | --- | --- | --- | --- | --- |
| One-way analysis of variance |  |  |  |  |  |
| P value | < 0.0001 |  |  |  |  |
| P value summary | *** |  |  |  |  |
| Are means signif. different? (P < 0.05) | Yes |  |  |  |  |
| Number of groups | 3 |  |  |  |  |
| F | 95.58 |  |  |  |  |
| R square | 0.8528 |  |  |  |  |
| Bartlett's test for equal variances |  |  |  |  |  |
| Bartlett's statistic (corrected) | 17.42 |  |  |  |  |
| P value | 0.0002 |  |  |  |  |
| P value summary | *** |  |  |  |  |
| Do the variances differ signif. (P < 0.05) | Yes |  |  |  |  |
| ANOVA Table | SS | df | MS |  |  |
| Treatment (between columns) | 2.714 | 2 | 1.357 |  |  |
| Residual (within columns) | 0.4685 | 33 | 0.01420 |  |  |
| Total | 3.182 | 35 |  |  |  |
| Bonferroni's Multiple Comparison Test | Mean Diff. | t | Significant?<br>P < 0.05? | Summary | 95% CI<br>of diff |
| 0 hrs of phage infection vs 2 hrs of phage infection | 0.4220 | 8.676 | Yes | *** | 0.3078 to<br>0.5362 |
| 0 hrs of phage infection vs 4 hrs of phage infection | 0.6645 | 13.66 | Yes | *** | 0.5503 to<br>0.7787 |

**Supplementary Figure 1.**
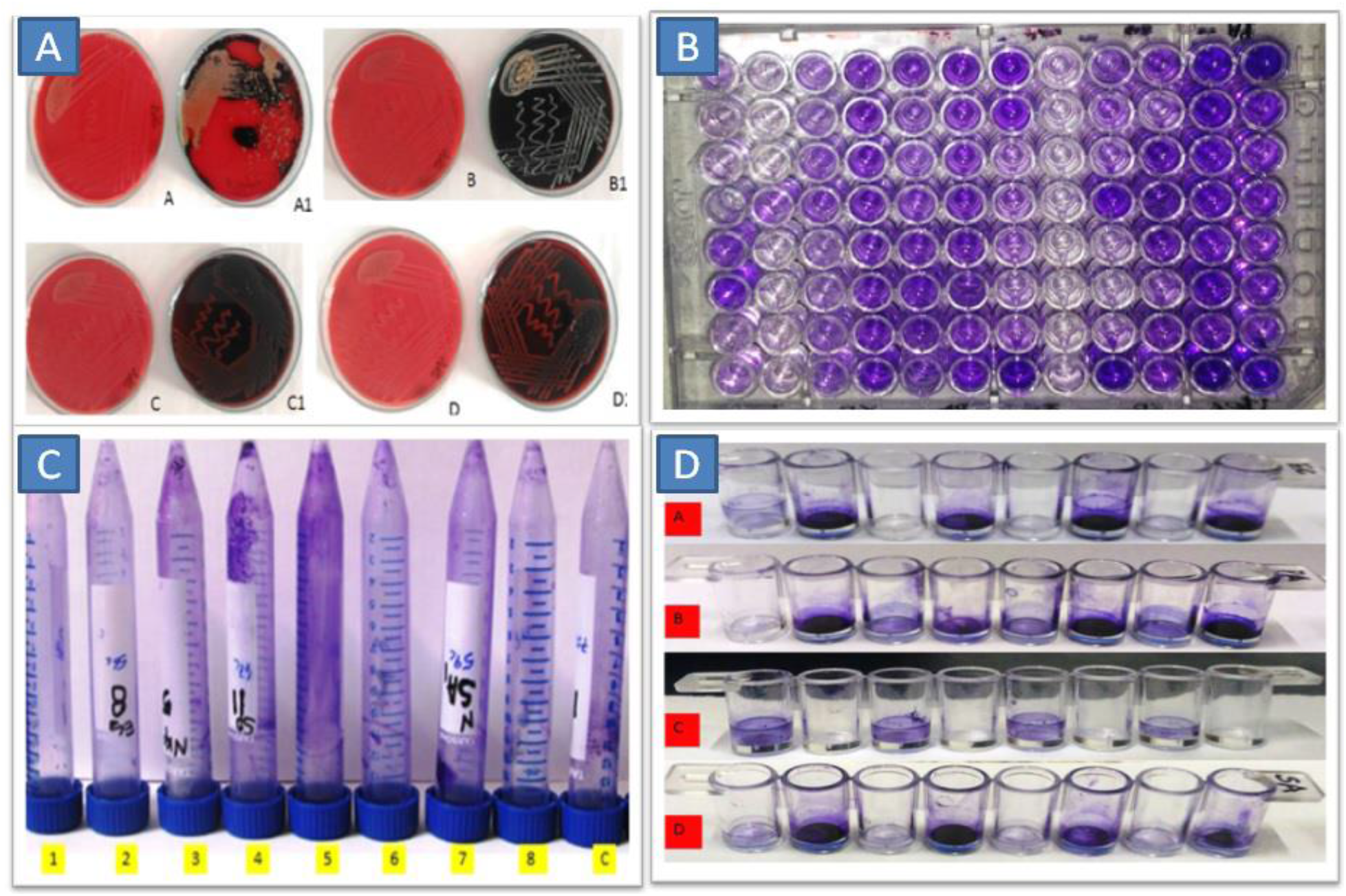
Qualitative detection of biofilm producers by Congo red agar, Microtiter plate and plastic test tube methods

**Supplementary Figure 2.**
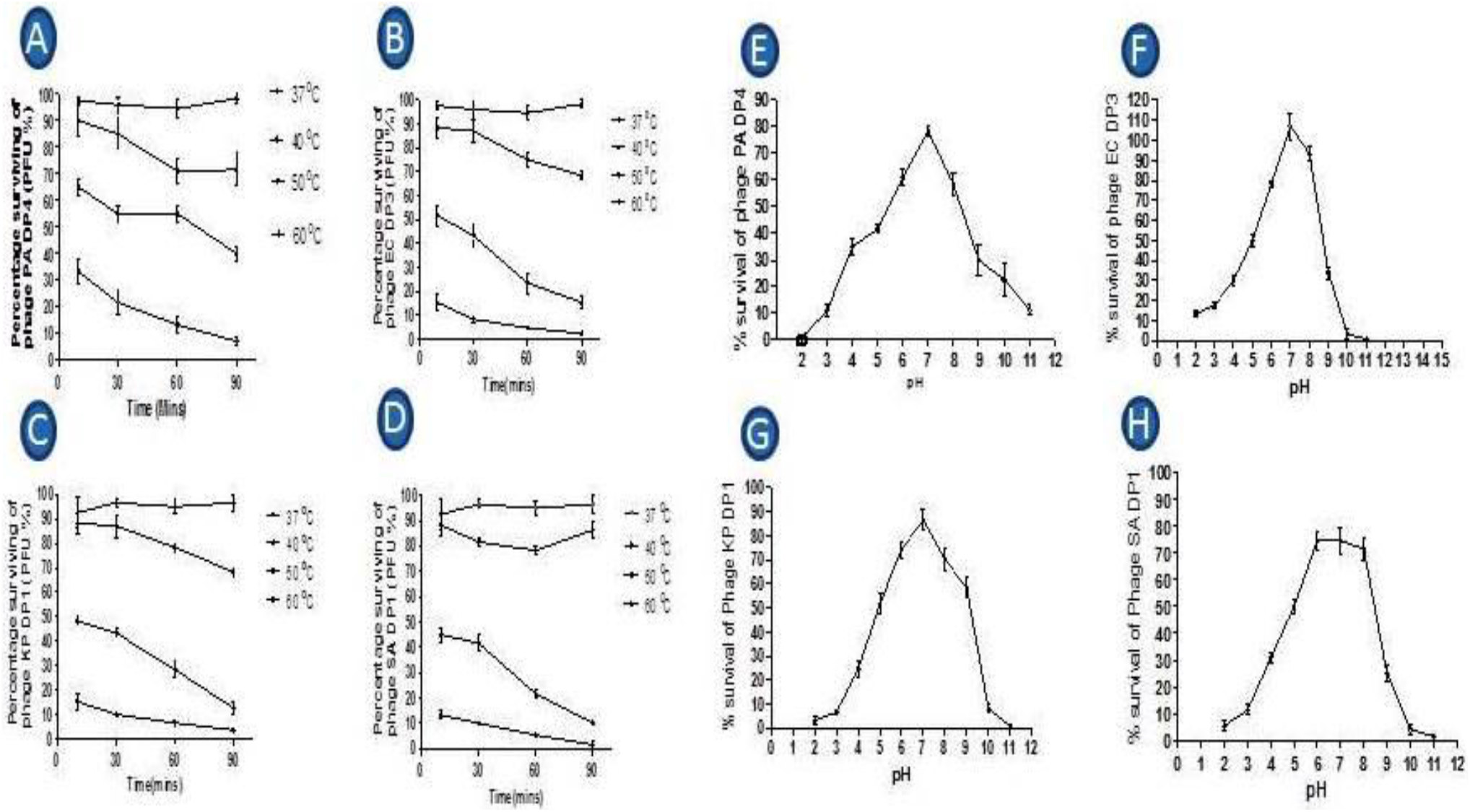
Effect of pH and Temperature on the bacteriophages vB_PAnP_PADP4, vB_SAnS_SADP1, vB_KPnM_KPDP1 and vB_ECnM_ECDP3. On the graphs, all values represented mean of three determinations ± SE by using Graph Pad Prism software.

